# Topological Learning Approach to Characterizing Biological Membranes

**DOI:** 10.1101/2023.11.28.569053

**Authors:** Andres S. Arango, Hyun Park, Emad Tajkhorshid

## Abstract

Biological membranes play key roles in cellular compartmentalization, structure, and its signaling pathways. At varying temperatures, individual membrane lipids sample from different configurations, a process that frequently leads to higher-order phase behavior and phenomena. Here we present a persistent homology-based method for quantifying the structural features of individual and bulk lipids, providing local and contextual information on lipid tail organization. Our method leverages the mathematical machinery of algebraic topology and machine learning to infer temperature-dependent structural information of lipids from static coordinates. To train our model, we generated multiple molecular dynamics trajectories of DPPC membranes at varying temperatures. A fingerprint was then constructed for each set of lipid coordinates by a persistent homology filtration, in which interactions spheres were grown around the lipid atoms while tracking their intersections. The sphere filtration formed a *simplicial complex* that captures enduring key *topological features* of the configuration landscape, using homology, yielding *persistence data*. Following fingerprint extraction for physiologically relevant temperatures, the persistence data were used to train an attention-based neural network for assignment of effective temperature values to selected membrane regions. Our persistence homology-based method captures the local structural effects, via effective temperature, of lipids adjacent to other membrane constituents, e.g. sterols and proteins. This topological learning approach can predict lipid effective temperatures from static coordinates across multiple spatial resolutions. The tool, called `MembTDA`, can be accessed at https://github.com/hyunp2/Memb-TDA.

## 1 Introduction

Biological membranes are crucial for cellular compartmentalization and structural integrity, as well as act a major platform for signaling pathways that govern environmental response.^1^ Membranes also serve as the primary boundary between pathogens and internal cellular compartments, thus being essential in both physiological and pathophysiological cellular response.^2,3^ Lipid membranes provide a structural context for functional protein conformations, for both peripheral and transmembrane proteins.^4–6^ Membrane composition is ubiquitously heterogeneous, consisting of variable lipids, sterols, and proteins.^7^ Individual lipids can sample from multiple acyl tail configurations, depending on local environment, pressure, and temperature, leading to distinct membrane properties.^8–10^

In the case of homogeneous lipid compositions, membranes experience temperature-dependent, higher order phase phenomena, for example, transformation between the gel and liquid phases, as a result of shifting lipid acyl tail configurational energy basins. Lipid order parameters (S_CH_/S_CD_) serve as a way to capture configurational properties of lipids.^11^ The order parameter for an acyl lipid tail is calculated using the angle, *θ*, formed between the bilayer normal and the carbon-hydrogen, or carbon-deuterium, bond vector:

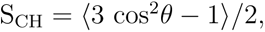

where the angular brackets, ⟨*…*⟩, represent temporal/molecular ensemble averages.^11^ Through molecular dynamics (MD) simulations, one can capture atomic representations of individual lipid configurations. When used in conjunction, order parameter calculations from MD trajectories can help refine force fields and provide bulk information on phase behavior. ^12^

Here we present a novel method for characterizing lipid order using a topological learning approach with MD, as an alternative to S_CD_ calculations. Our approach learns the underlying temperature-dependent, potential energy surface from a wide range of lipid configurations, obeying a Boltzmann distribution sampled from equilibrium MD simulations, providing an effective temperature estimate for individual lipids (Figure 1).

**Figure 1:**
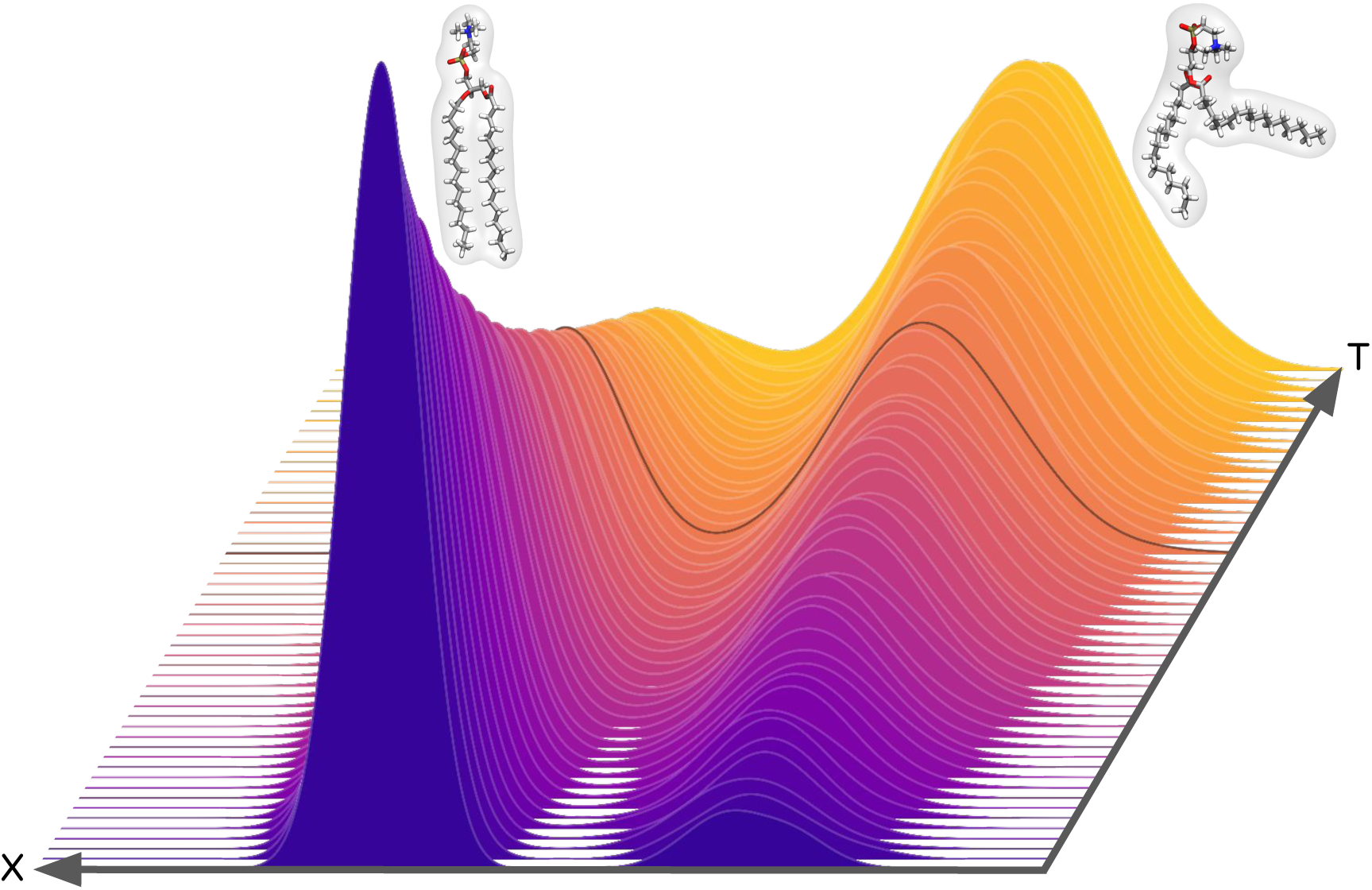
Schematic view of the configurational manifold mapped by `MembTDA`. Here we show configuration space X versus MD input temperatures T. The height represents a configuration probability density, with the phase transition temperature shown as a black line where two prominent configurations are of equal probability. `MembTDA` maps out this surface and infers a distribution of likely configurational temperatures, based on potential energy features derived from static coordinates, for which an expectation value yields an effective temperature.

Our method, named `MembTDA`, leverages topology, a branch of mathematics concerning sets that contain an inherent structure that is preserved under continuous deformations. Topological data analysis (TDA)^13^ has rapidly become one of the main tools used by artificial intelligence researchers to agnostically extract feature information from various data sets. TDA focuses on the inherent topological features of the data, to create a topological fingerprint represented as bar codes, diagrams, or images (Figure 2). One clear benefit of TDA is the ability to cluster data consistently over other convergence-based methods, as well as being resilient to perturbations, specifically with the use of TDA techniques like persistent homology (PH).^14,15^ PH is a method of extracting both geometric and topological information from a point cloud, like atomic coordinates. Major benefits of the PH methodology include robustness under perturbations, characterization of points clouds of varying densities, and the potential for abstract feature comparison via the Wasserstein distance.^14–16^ The robustness of PH lends itself nicely to the complex nature of biological data analysis. In the case of biological membranes, lipids can have variable acyl tails as well as variable headgroups;^17^ these variable factors and underlying constant features can be captured using PH even under perturbative effects, like dynamics. Capturing conserved topological features of membrane data using persistence homology, allows us to train neural networks for biophysical and thermodynamic feature prediction, like temperature.

**Figure 2:**
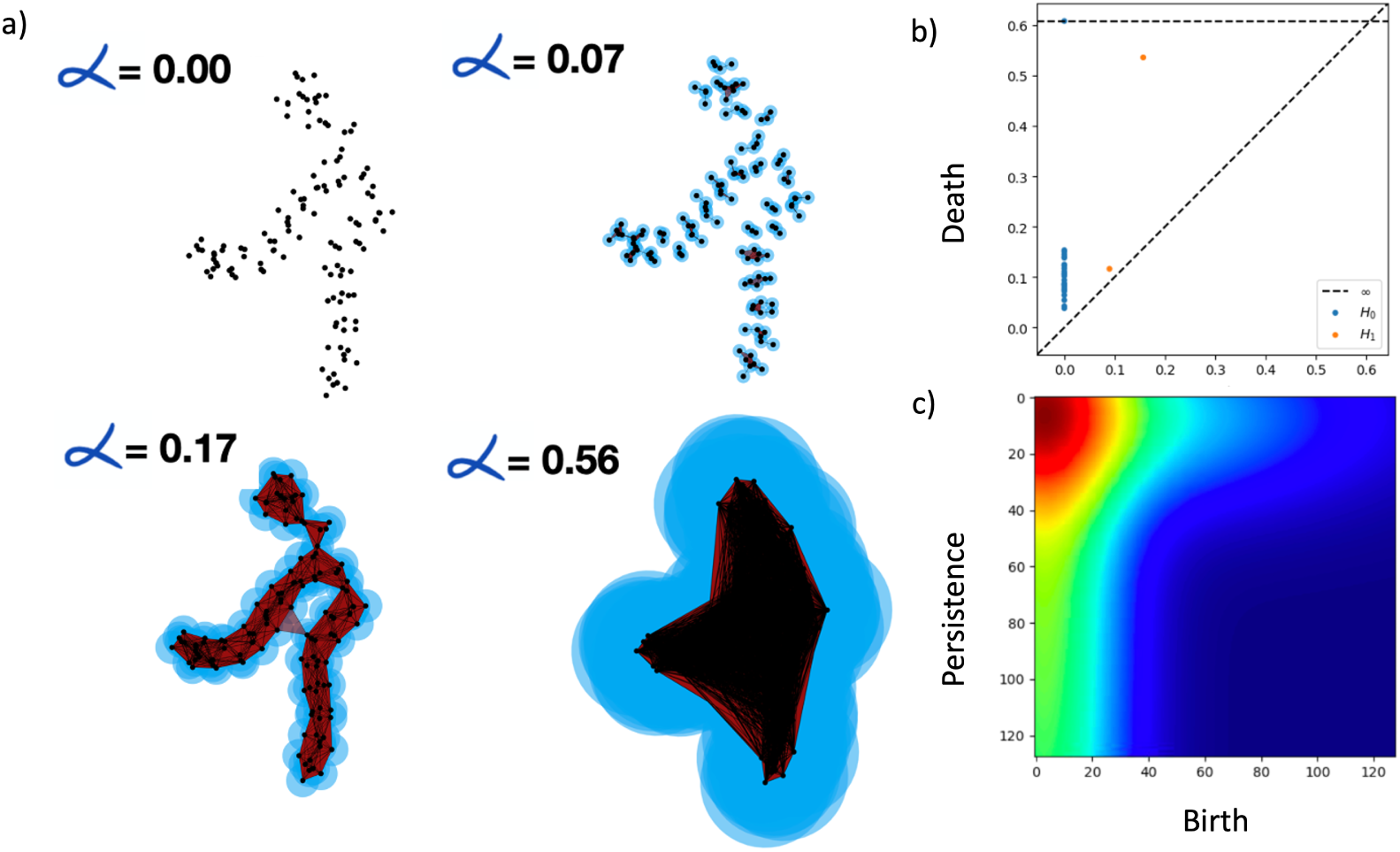
Overview of persistent homology (PH) calculations. a) To extract topological features via PH, we first construct a simplicial complex on the point clouds obtained from lipid atomic coordinates. A filtration value, *α*, is increased, which increases the radii of spheres surrounding every point in the point cloud. An example simplicial complex at varying filtration values is shown. b) The topological features in the simplicial complex can be represented as a persistence diagram depicting birth versus death of features. c) Plotting persistence (death minus birth) versus birth, and performing a Gaussian kernel approximation on the diagram yield an image that is amenable to visual transformers, called a persistence image.

Machine learning (ML) or deep neural network can be used to extract hidden patterns that are usually hard to detect using conventional techniques. These include, but not limited to, predicting protein structures to predicting qualitative or quantitative molecular properties.^18–23^ In this work we train a deep neural network model specialized in processing images, such as Visual Transformer (`ViT`)^24^ ^25^ or `ConvNeXt`,^26^ with persistent data information of lipid coordinates at varying temperatures. By marrying ML with TDA, we demonstrate our model’s capability in predicting individual lipid’s effective temperatures. `MembTDA` maps out the configurational manifold of lipids and allows for inferring an effective temperature from a distribution of likely configurational temperatures Figure 1.

## 2 Methods

Our approach is comprised of three major elements. First, we create a molecular data set for lipids at varying temperatures using molecular dynamics (MD) simulations. Then we featurize the MD data using PH (specifically, persistence images). Lastly, we train an attention based transformer using the data from PH and MD.

### 2.1 Molecular Dynamics Simulations

A major part of model development is data accumulation. Our data set consisted of trajectories from MD simulations of membranes in 51 different temperatures ranging 280–330 K, spaced apart by 1 K. The model lipid bilayer consisted of 117 dipalmitoyl-phosphatidylcholine (DPPC) lipids per leaflet and was constructed using the CHARMM-GUI webserver.^27^ The system was solvated and ionized with 150 mM NaCl to mimic cellular conditions. The CHARMM36m^28^ force field parameters and the TIP3P water model^29^ were used for all the simulations. The equilibration of the systems was performed using the NAMD.^30,31^ A 100-ns production run was then performed for each temperature replica with GPU-resident NAMD to ensure optimal GPU scaling.^31^ Observed gel and liquid phase transitions in the simulated bilayers were captured within the first 20 ns of each simulation replica. Only the last 200 frames (2 ns) were used for model training to minimize the effects of the degenerate starting conditions. In total, the simulation sampling amounted to 100 ns per replica, approximately an aggregate of 5 *µ*s sampling.

As in the equilibration runs, the production simulations were performed as an NPT ensemble at varying temperatures from 280 K to 330 K with a pressure of 1.0 atm. An integration time step of 2 fs was used throughout. The Nośe-Hoover Langevin piston method^32–34^ was used to maintain a constant pressure, with temperature maintained constant via Langevin dynamics with a 0.5 *ps*^−1^ damping coefficient.^35,36^ A 12-Å cutoff was used for nonbonded interactions with a smoothing function implemented after 10 Å. The bond distances of the hydrogen atoms were constrained using the SHAKE algorithm.^37^ For long-range electrostatic calculations, the particle mesh Ewald (PME) method^38^ was used, with a grid density of more than 1 Å^−3^. Visualization and analyses of the simulations were done using Visual Molecular Dynamics (VMD)^39^ and MDAnalysis.^40^

### 2.2 Persistent Homology

Persistent homology (PH) is a method of extracting both geometric and topological information from a simplicial complex constructed from a point cloud, like atomic coordinates. A simplicial complex K is a collection of simplices in R*^n^*. Simplices are topological descriptions of connected points, where a single point is a 0-simplex, two connected points form a 1-simplex, three points a 2-simplex, and so on (Figure 2). In the case of a point cloud, like the coordinates of a phospholipid bilayer or a protein, we have a collection of 0-simplices, i.e., isolated points.

In PH, we measure the unique fingerprint of a point cloud (atomic coordinates in the case of lipids) by tracking the topology of its simplicial complex at a varying filtration parameter *α*. To vary the filter parameter, we grow *n*-dimensional spheres around each vertex (individual point) of the point cloud. In our work, *n* = 3, because the points represent the atomic Cartesian coordinates of lipids. As the radius, the filtration parameter *α*, of the spheres increases, the ratios of *n*-simplices vary depending on the intrinsic structure of the data. We perform a filtration of the 0-simplicial complex by growing *n*-dimensional spheres around each point, starting with a radius of *α* = 0 and fully disconnected 0-simplices. As we increase *α*, we keep track of overlaps between neighboring spheres such that two overlapping spheres form a 1-simplex (i.e., an edge), trisecting spheres form a 2-simplex (i.e., a triangle face), and so on.

More symbolically, starting from a point cloud {*x*_1_*, x*_2_*, …, x_n_*}, as in the case of the Cartesian coordinates, we can construct a simplicial complex by growing *n*-dimensional spheres around each point and tracking sphere intersections. In the case of an *α*-complex construction,^41^ we begin by creating a Voronoi partition, *V_x_*, of our point cloud:

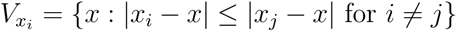

Using the condition that our *α*-complex can only reside within our Voronoi partition, we then grow spheres, *B_α_*, around each point:

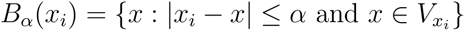

The process of growing spheres around each vertex is referred to as a filtration. Taken together, the intersection of our simplices at every radius *α* forms a simplicial complex, in this case an *α*-complex:

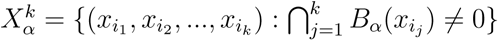

Our *α*-complex, *X_α_^k^*, contains information of the connectivity of our system at varying radii, *α*. We then identify holes in our data by applying homology to the *α*-complex.^42^

Ultimately, the homology of the varying filtration value *α* yields information on topological and geometric features. We can visualize the birth and death of the captured features obtained from PH from the varying filtration value using persistence barcodes, persistence diagrams, and persistence images, all of which are examples of persistence data Figure 2-a. Applying *homology* to the simplicial/*α*-complex, comprised of the simplices at all varying radii, yields topological features such as birth and death information of 0-, 1-, 2- homology groups (i.e., connected components, holes, and voids); we denote these homology groups *H*_0_*, H*_1_ and *H*_2_, respectively. The topological features attained from PH are primarily represented visually as persistence diagrams and persistence images (Figure 2-b, c). In our work, we interchangeably use terms PH, persistence data, topological fingerprints, or topological features, to refer to extracted topological data obtained from the TDA approach.

Although persistence barcodes and persistence diagrams are readily interpretable for humans, their sparse nature is less amenable to learn patterns for deep learning-based models such as visual transformers (`ViT`)^24^ using the attention mechanism^43^ or convolutional neural networks. To address the sparse nature of persistence diagrams, we transform them into persistence images, ^42^ a process called PH vectorization, which leverages Gaussian kernel approximations to create topological fingerprints that can be readily fed through a computer vision based deep neural network to learn complex patterns and predict properties. The reasons for choosing vision based deep neural network architectures are further elaborated in subsection 2.3.

Instead of birth versus death information, persistence images have persistence versus birth; where persistence is characterized by death minus birth.^42^ Using every 10 ps of the last 2 ns of the membrane simulations at variable temperatures, coordinate data of individual lipids were used to form the persistence images Figure 3.

**Figure 3:**
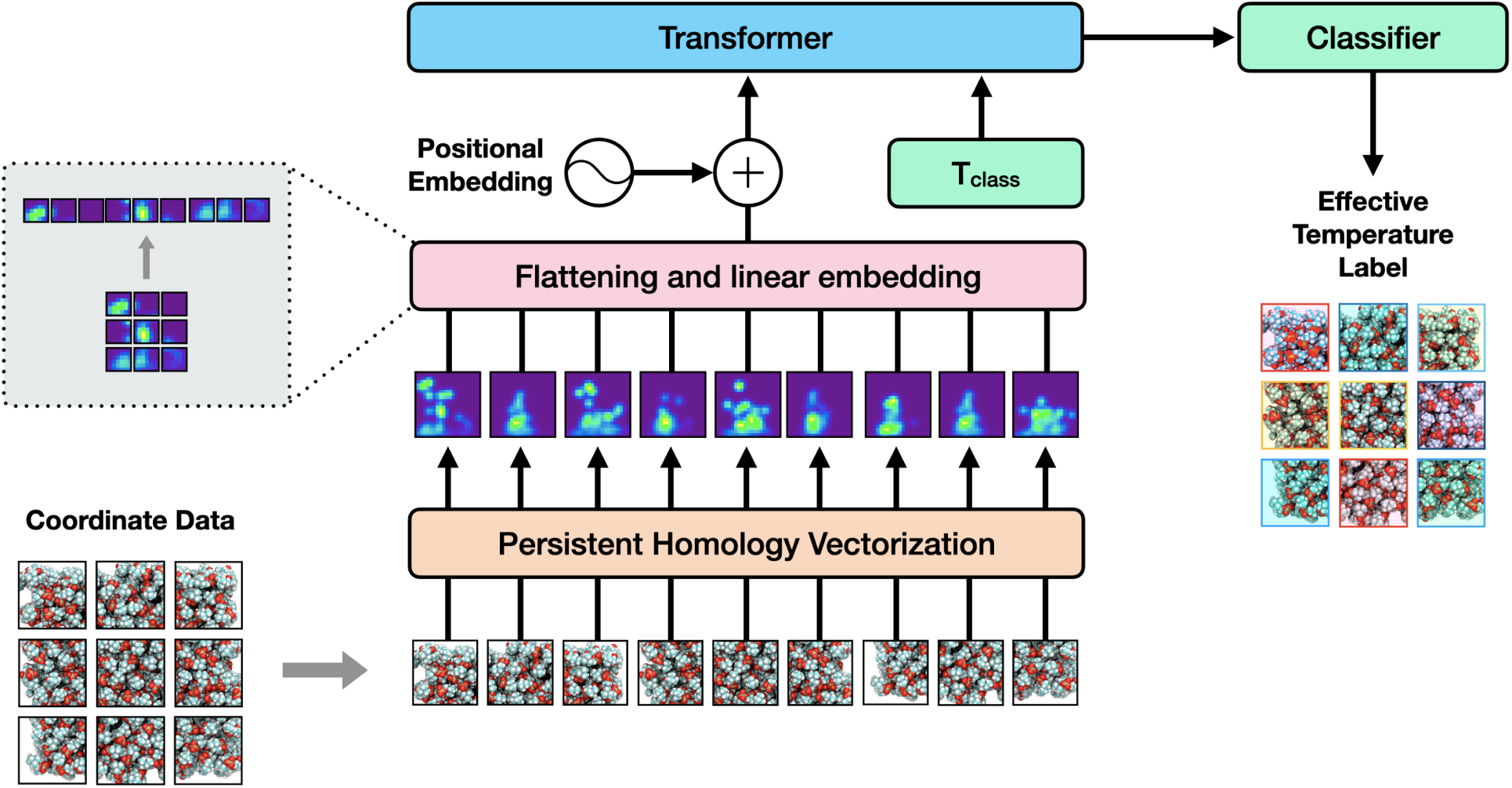
`MembTDA` overall workflow. First MD data are fed as an input to the workflow as coordinate patches, either as individual lipids or as patches. The atomic coordinates then undergo PH vectorization (diagram to image transformation), in which topological features are characterized and represented as persistence images. Each persistence image undergoes a flattening and linear embedding step, including patching and positional embedding. The embedded data are then fed through a transformer (`ViT`), with a temperature class label from the MD input temperature. The model ultimately acts as a classifier with a distribution of probability values for possible temperature classes, for which an expected value yields an effective temperature *T*_E_.

### 2.3 Deep Learning Model for Processing Persistence Images

In our work, we used `ViT` architecture for prediction of effective temperatures. The `ViT` architecture we used was based on window attention of `Swin Transformer version 2`^25^.^44^ The objective function is the temperature class prediction, *L*_CE_(*p_θ_*(*T_i_*)*, T*_true_), and expected temperature prediction, *L*_MSE_(**E***_i_, T*_true_). By training `MembTDA` with bi-objective functions, we can optimize our neural network model for a more robust representation learning of our input. Our overall workflow is described in Figure 3 and elaborated further in Figure 5.

## 3 Results

Lipid bilayer phase transitions are readily characterized experimentally by metrics such as heat capacity and acyl tail order parameters. Our topological deep learning approach for characterizing lipids allows us to probe lipid phases for individual lipid molecules. We demonstrate the utility of `MembTDA` on homogeneous and heterogeneous membranes, including, lipid bilayers simulated at variable temperatures, and bilayers with transmembrane or peripheral protein systems.

### 3.1 Homogeneous Membranes at Variable Temperatures

To assess the ability of `MembTDA` to identify and classify lipid phases, we performed inference on the lipid training set and report the effective temperature distribution in Figure 4-b. Inference on the initial training set’s effective temperatures, the distribution obtained by calculating expected values of `MembTDA` output classes, reveal a seemingly bimodal distribution with internal effective temperature minima at 297.29 K and 306.44 K.

**Figure 4:**
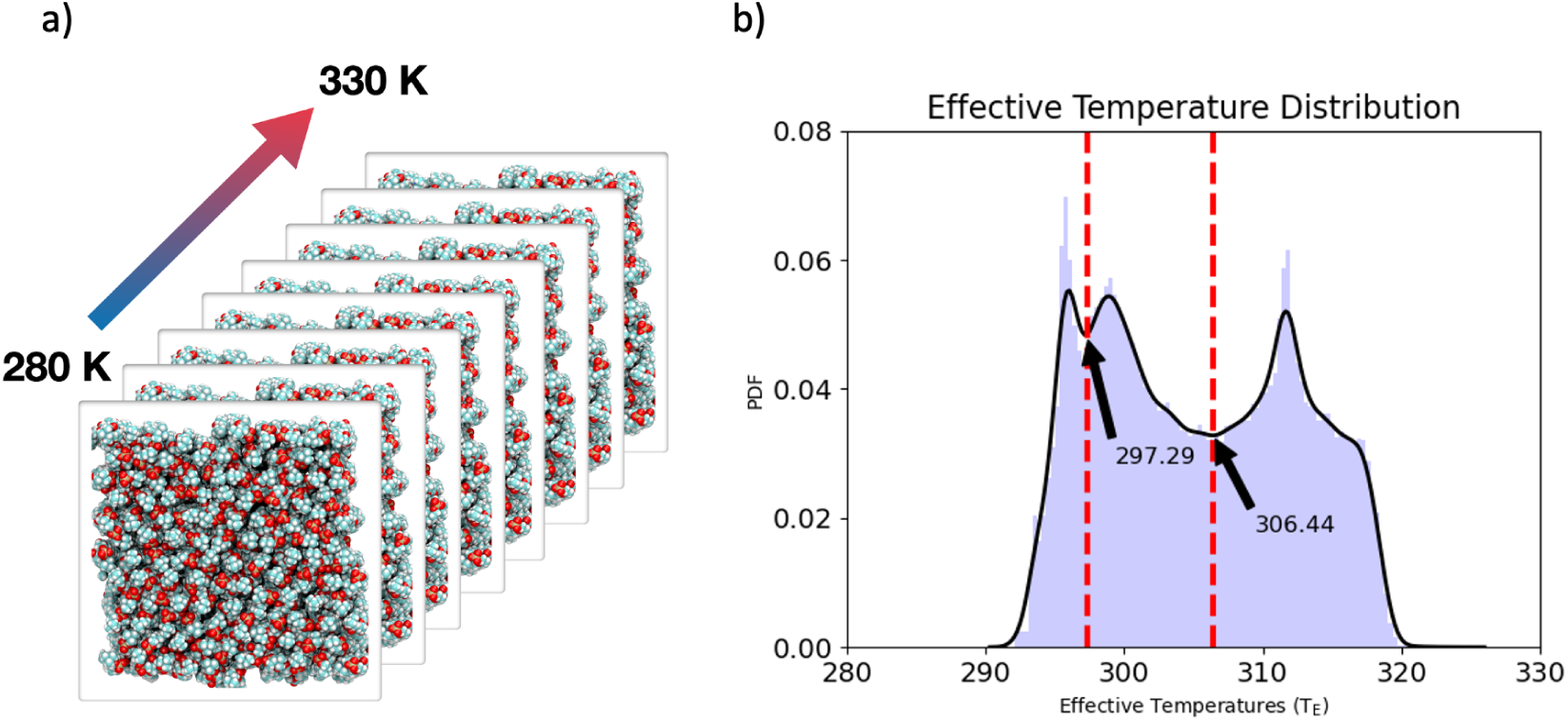
Inference of entire MD data set. a) A small subset of DPPC lipids from membrane simulations across 51 temperatures (280-330 K) was used to train the `ViT`-based `MembTDA`. b) Applying the trained `MembTDA` back on the training set to capture temperature class expectation values yielded a bimodal effective temperature distribution. `MembTDA` effectively classifies lipids as either gel or liquid phase. `MembTDA` also captures a *L_α_*/*L_β_* phase transition at 306.44 K, and acyl tail tilting at 297.29 K.

According to Khakbaz et. al.,^45^ 308.15 K is where *L_α_* (crystalline liquid) to *L_β_/P_β_* (gel/ripple, respectively) transition occurs, suggesting a sharp decrease in surface area at this temperature. In addition, according to the authors,^45^ at 298.15 K, the *L_β_* phase occurs with predominantly tilted acyl chains with respect to the membrane normal. Our MD simulation environment closely matched that used in Khakbaz et. al.^45^ (i.e., NAMD,^30,31^ Charmm36^46^ parameters, DPPC lipids), from which `MembTDA` predicted two biophysically significant temperatures, purely from persistence data followed by neural network operation. The results demonstrate that `MembTDA` is robust at capturing this aspect of the lipid membranes, in this case phase transitions. The absolute errors between temperatures from Khakbaz et. al.^45^ and `MembTDA` predictions of melting and chain tilting temperature is 1.7 K and 0.86 K, respectively.

### 3.2 Transmembrane protein in POPC – AQP5

Aquaporin 5 (AQP5), a protein involved in water homeostasis, was previously simulated in our lab in a 1-palmitoyl-2-oleoyl-*sn*-glycerol-3-phosphocholine (POPC) bilayer with a thermostat input temperature of 310 K. Unlike DPPC, POPC contains asymmetrical acyl tail lengths, altering its melting temperature and membrane dynamics. ^47^ Inference on this system revealed an effective temperature distribution above the melting temperature of DPPC and POPC, indicative of highly disordered lipids, potentially due to protein-lipid interactions.

### 3.3 Transmembrane protein in POPE – LaINDY

LaINDY, a transmembrane bacterial transporter, was simulated previously in our lab in a 1-palmitoyl-2-oleoyl-*sn*-glycero-3-phosphoethanolamine (POPE) membrane. Like POPC, POPE contains asymmetrical acyl tails, but also has a different head group (ethanolamine) than both DPPC and POPC. Inference on this system revealed two distinct peaks, potentially indicative of variable protein-lipid interactions between LaINDY and POPE. This analysis demonstrates the ability of `MembTDA` to perform inference on lipid head groups outside the training data.

### 3.4 Peripheral Membrane protein in POPC/POPS - *β*2GP1

*β*2GP1 a peripheral membrane protein known to bind 1-palmitoyl-2-oleoyl-*sn*-glycero-3- phospho-L-serine (POPS) lipids, was simulated previously in our lab in an equal ratio of POPC:POPS membrane at 310 K. Effective temperature inference on this heterogeneous membrane revealed two distinct distributions across the membrane. Upon decomposition of POPS and POPC effective temperatures, we identified that `MembTDA` is able to distinguish between POPC and POPS lipids. We find this remarkable since our model was trained purely on DPPC persistence data, and the input point cloud contained no information on charge, atom types, or connectivity. Furthermore, the low effective temperature distribution captured for POPS is interesting in the context of *β*2GP1 binding specificity, since *β*2GP1 is known to selectively bind POPS-rich membranes. Furthermore, we observe that the distribution of effective temperatures for POPC from the decomposed heterogeneous membranes, closely resembles the distribution observed for AQP5-embedded POPC.

In the Supplemental Figure S11 and Figure S12, we show the overall *H*_1_ Wasserstein matching diagram and close-up view of the same diagram in the 0.2-0.4 birth-death range for a better demonstration. We chose 100 random lipids, each from three temperature categories, i.e. low, melting, and high temperatures, resulting in a total of 300 samples, calculated persistence diagrams and overlaid them with colors labeled in the legend (e.g., labeled “all lower temps” with an “o” marker). Then we calculate barycenters of each temperature categories 100 sample persistence diagram birth-death points (e.g. labeled ”lower temp”, with an ”X” marker). Barycenters are centroids of non-linear systems such as birth-death points of persistence diagrams.^48^ The Wasserstein distance is a metric to compare the similarity between two persistence diagrams. The calculation of the Wasserstein distance discerns which points of one diagram are similar to those of another diagram, hence *matching* birth-death points between two diagrams. These *matching* points are represented as an edge connecting two barycenter points in supplemental Figure S12 (i.e. blue ”X” to yellow ”X” and to red ”X”).

From the Wasserstein diagrams, we can conclude that there are prominent *H*_1_ features (holes) appearing and persisting about 0.2 *α* radius. When observed closer (supplemental Figure S12), we can see that barycenters of three different temperature categories persistence diagrams show movements, indicated by edge connections. This shows that there distinct persistence data encoded across three lipid temperature categories, indicating a temperature dependent flow of topological features. This is consistent with the idea that `MembTDA` maps out the underlying configurational landscape as seen in Figure 1.

## 4 Discussion

We present a novel lipid characterization method, termed `MembTDA`, that estimates effective temperature of lipids from static coordinates, as a topological alternative to the commonly used *S_CH_/S_CD_* order parameter calculations. We demonstrate `MembTDA`’s effectiveness as a functional classifier by performing inference on homogeneous and heterogeneous membranes, with variable tails (e.g., DPPC, POPC), head groups (e.g., POPC, POPE, and POPS), and in the presence of proteins (e.g., AQP5, LaINDY, and *β*2GP1). Although our methodology functionally acts as a classifier, the reliance on coordinate data, i.e., point cloud information, necessarily implies that the method is inherently based on potential energy features, which are functions of *xyz*-coordinates. Furthermore, since temperature is predominately calculated as a kinetic energy parameter from the equipartition theorem,^49^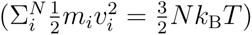, the effective temperatures of static snapshots we report are not exactly representative of traditional bulk temperature, an *ensemble* property. The effective temperature we report is a quantity obtained by taking the expected value of the `MembTDA` output distribution, which is based on a mapping of the configurational landscape with an associated input MD temperature (Figure 4).

More accurate predictions of the local temperature may be possible by taking velocity information from simulation, and using the equipartition theorem, 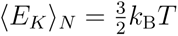, to estimate individual lipid temperatures. Typically the average kinetic energy of the entire simulation system, including all non-lipid constituents, is used to calculate the system’s temperature:

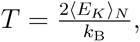

accounting for degrees of freedom N as 3N-3 (Supplemental Figure S2, Supplemental Figure S3, Supplemental Figure S4). Due to the variable molecular degrees of freedom arising from differing intra-molecular (including the number of atoms, bonds, angles, and dihedrals) and inter-molecular interactions between lipids (contextual information such as lipid-lipid or lipid-protein association), the use of the equipartition theorem to estimate individual lipid temperatures based on their kinetic energy is non-trivial.^50^ Estimates of individual lipid temperatures using kinetic energy formulations of temperature yield individual lipid temperatures, sometimes over 50 K below the global system temperature (Supplemental Figure S5, Supplemental Figure S6, Supplemental Figure S7). Alternatively to a kinetic energy-based estimate for temperature, a configurational temperature may be also calculated using the Jepps formulation:

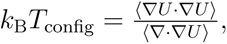

where U is the system’s potential energy and ⟨*…*⟩ denotes an ensemble average.^51^ Although based on the potential energy, calculation of the configurational temperature from the Jepps formulation ultimately requires computationally expensive Hessian calculations. The term ⟨∇ · ∇*U* ⟩ necessitates calculating the divergence of the gradient of the potential energy, which requires a Hessian and degrees of freedom information, making accurate configurational estimates prohibitive. ^51,52^ Furthermore, in practice, MD users primarily only store atomic coordinates to save on storage and computation costs; only writing out velocity information for restart purposes.^53^ `MembTDA` provides a pragmatic way to estimate effective temperatures from data typically saved by MD practitioners, allowing post-processing of extant simulation trajectories. In addition, the ability to analyze static coordinates lends to potential applications in analyzing effective temperatures in novel structural data.

`MembTDA` was originally trained on DPPC membrane structures across different temperatures. We demonstrate that the so trained `MembTDA` can capture two important temperatures of DPPC membranes, namely chain tilting and melting temperatures as mentioned in sub- section 3.1. This result is particularly interesting since `MembTDA` was only given persistence data to classify into one of the 51 temperature classes, and was trained only on a partial subset. However, once trained, when given all the lipid tail dataset, `MembTDA` predicts critical unique temperatures that were never explicitly set as a training objective or part of the neural network architecture. The information captured in the persistence data and learned by the neural network has shown that the two local minima of the DPPC effective temperature distribution curve are indeed biophysically relevant temperatures as reported in Khakbaz et al.^45^

The effective temperatures predicted by `MembTDA` are dynamics of lipids, since temperature prediction inherently takes velocity information^49^ into account. Also, for the configurational temperature, as in Jepps et al.,^51^ force information is taken into account. Both force and velocity are vectors projected on the atomic position, responsible for propagating the MD integration. However, since there are no direct atom velocity nor complete force information (due to individual lipid force being considered without environment) in static lipid coordinates, we can speculate that persistence data carry mixed information of both velocity (a proxy for *entropic* information)^54^ and force information for which the neural network is effectively able to learn. Such hidden information comes in the form of persistence data, implying that effective temperature information can be retrieved by accounting for connected components and holes present in the lipid configurations. To us, this implies that `MembTDA` maps the underlying configurational manifold and its topology, which is necessary for its physical dynamics (Figure 1).

Moreover, we have shown that `MembTDA` can capture effective temperatures of lipids in membranes with different lipid compositions other than what was used in its training, and for lipids in the presence of proteins. The dynamics of lipids are shown in Figure 5 where higher effective temperature lipids, colored in red, can be observed near the periphery of proteins. Also, different lipid types (i.e., POPC, POPE, and POPS) experience different effective temperatures because different head groups have different favorability to the protein residues they are interacting with, due to polarity and charges based interactions (an *enthalpic* effect), altering allowed configurational states.

**Figure 5:**
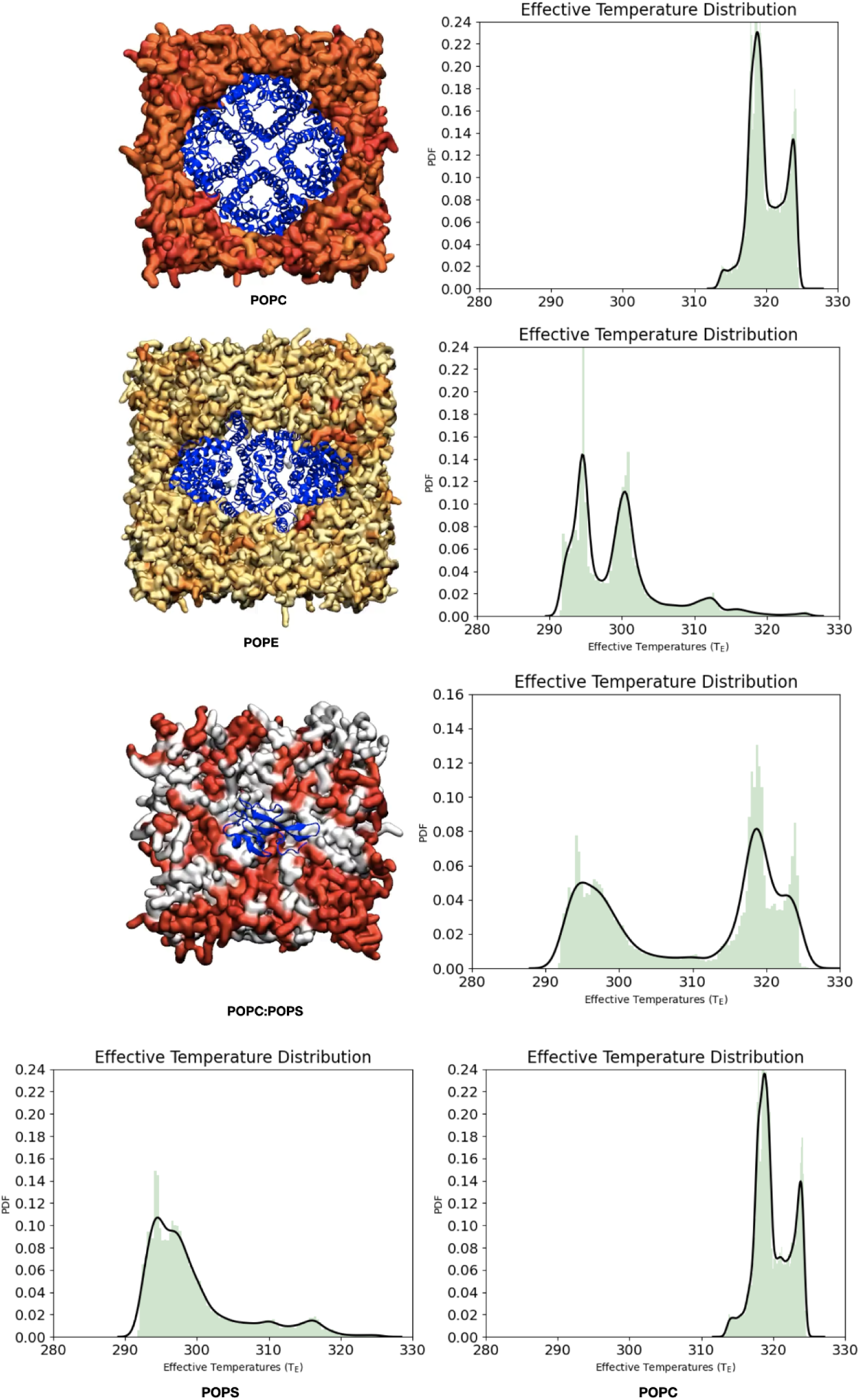
Inference on out of training data distribution. Effective temperature estimates for heterogeneous membranes containing varying proteins.

As for why `MembTDA` has accurate effective temperature predictability, we ascribe this to attention maps presented in Supplemental Figure S8, Figure S9, and Figure S10. For low-temperature lipids, we chose 16 random (hence 4 x 4 panels) lipids and extracted the attention maps `MembTDA` focuses on. An attention map (in our case, GradCAM ^55^) is an explainable AI (XAI) technique to highlight which features of the input an ML model focuses on to make a prediction. We can see that there is a semi-circle on the top left part of the persistence image data, which `MembTDA` identifies as important features (Supplemental Figure S8). The semi-circle indicates prominent *H*_1_ features (i.e., holes) in the persistence data which may be born at relatively earlier *α*-filtration values. Since lower temperatures render accessible lipid configurations to more rigid states, the attention map (GradCAM) can capture hole patterns more readily. On the other hand, for high-temperature lipids, the attention map (GradCAM) does not have a distinct pattern captured by our neural network (Supplemental Figure S9). This implies that a high variability of lipid configurations exists at high temperatures, and thus no consolidated patterns like those observed for low-temperature lipids. As for the melting temperature (Supplemental Figure S10), we see mixed patterns where nearly half the attention maps (GradCAM) resemble those of low-temperature lipids (i.e., prominent attention patterns), and the rest resemble those of high-temperature lipids (i.e., no particular attention patterns). We postulate that this is due to expected equal fractions of *L_β_* and *L_α_* phases at the melting temperature.

At all scales, entropic contributions create system heterogeneity that result in local order.^56^ In the context of biological membranes, we have shown that local order plays a role in lipid dynamics, by capturing local effective temperatures of individual lipids. The ability to recapitulate DPPC melting temperatures, demonstrated by `MembTDA`, reveals that PH is likely correlated to physical phenomena, potentially via a mapping of a manifold embedding representative of a potential energy surface with an inherent characteristic topology. We speculate that the effectiveness of our topological learning approach implies the possibility of an underlying analytical framework suitable for estimating physical properties such as melting temperatures or heat capacity, like a hybrid of the equipartition theorem and the Jepps formulation. More precisely, we posit that the use of PH on atomic data captures inherent topological features of the potential energy surface as shown in Figure 1 and Figure 4.

## Acknowledgements

We would like to thank Dr. Pochao Wen, Dr. Archit Vasan, Ali Rasouli, and Matt Sinclair for their feedback and useful commentary. The authors acknowledge support from the National Institute of General Medical Sciences of the National Institutes of Health under awards P41-GM104601, R24-GM145965, and R01-GM123455. ASA would like to acknowledge a graduate student fellowship from xxxxx. National Institutes of Health under award F31-HL136155. The content is solely the responsibility of the authors and does not necessarily represent the official views of the National Institutes of Health.

