## Supplemental Information for "Topological Learning Approach to Characterizing Biological Membranes"

### 5 Supplemental Materials

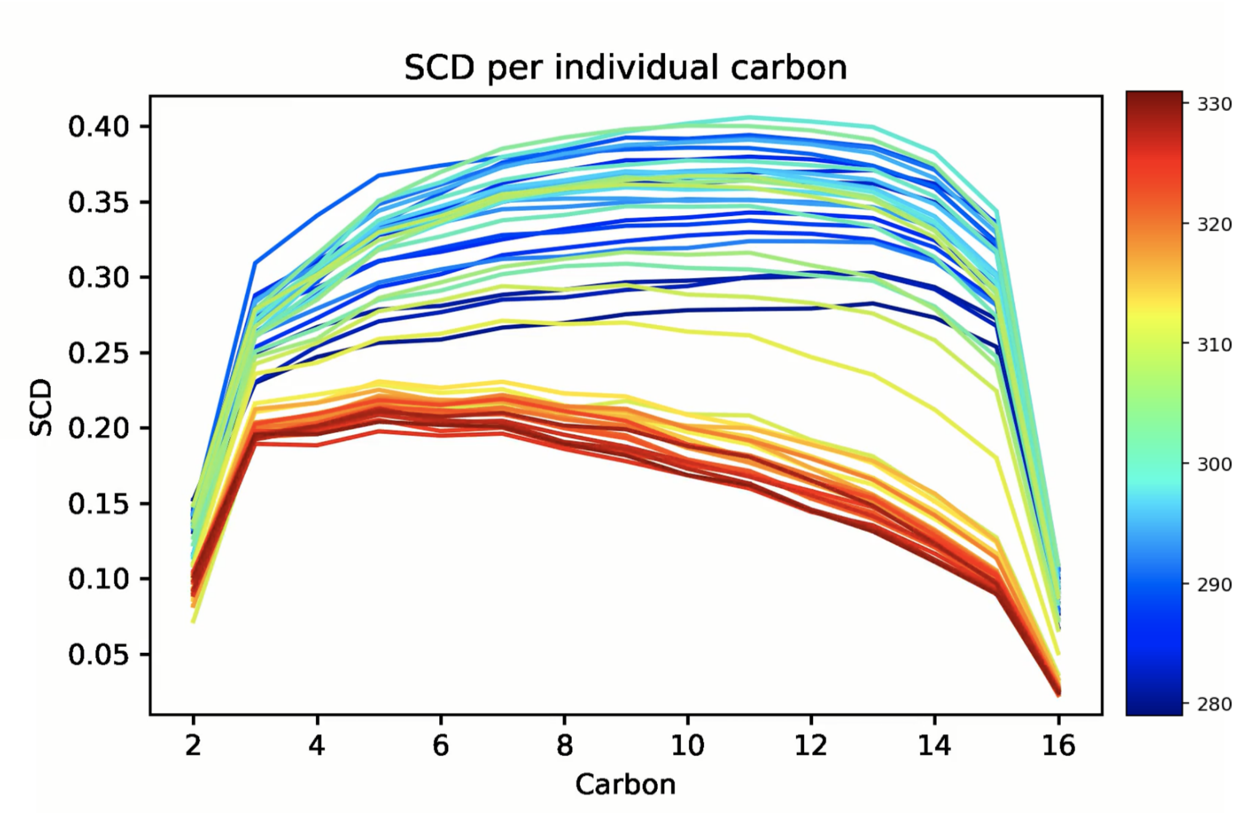


Figure S1: Ensemble average order parameter ($y$-axis) for DPPC lipid acyl tail carbons ($x$-axis) at different temperatures (color bar).

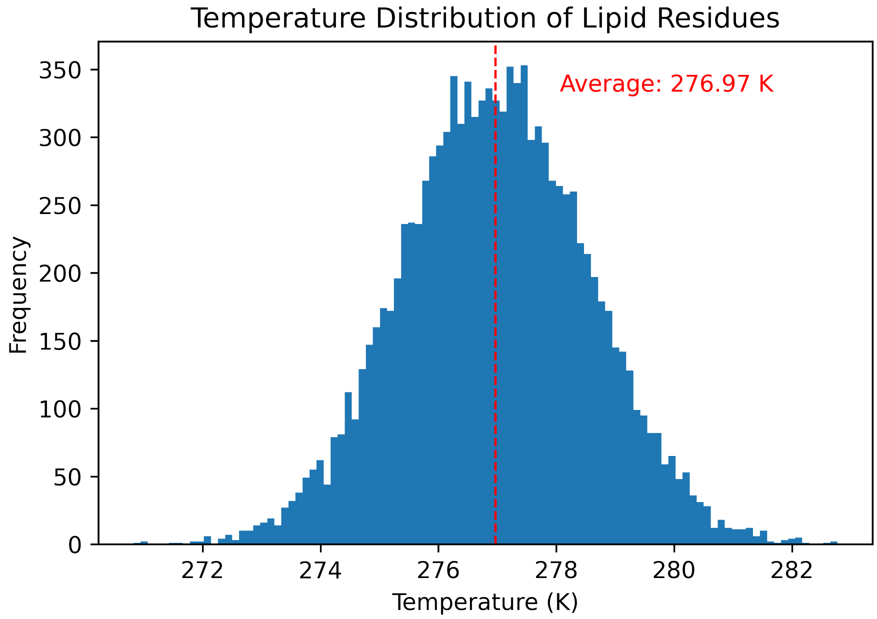


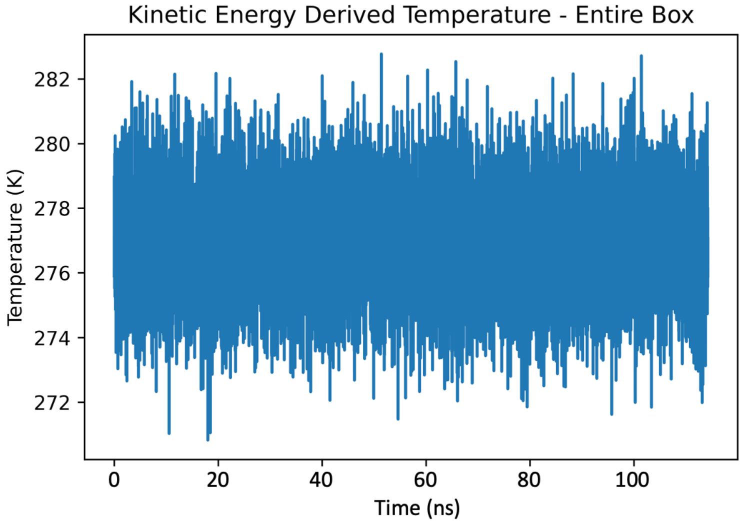


Figure S2: Top: Distribution of the kinetic energy-based temperatures, calculated with the equipartition theorem using an MD input temperature of 280 K. Bottom: Time series of the kinetic energy based temperatures.

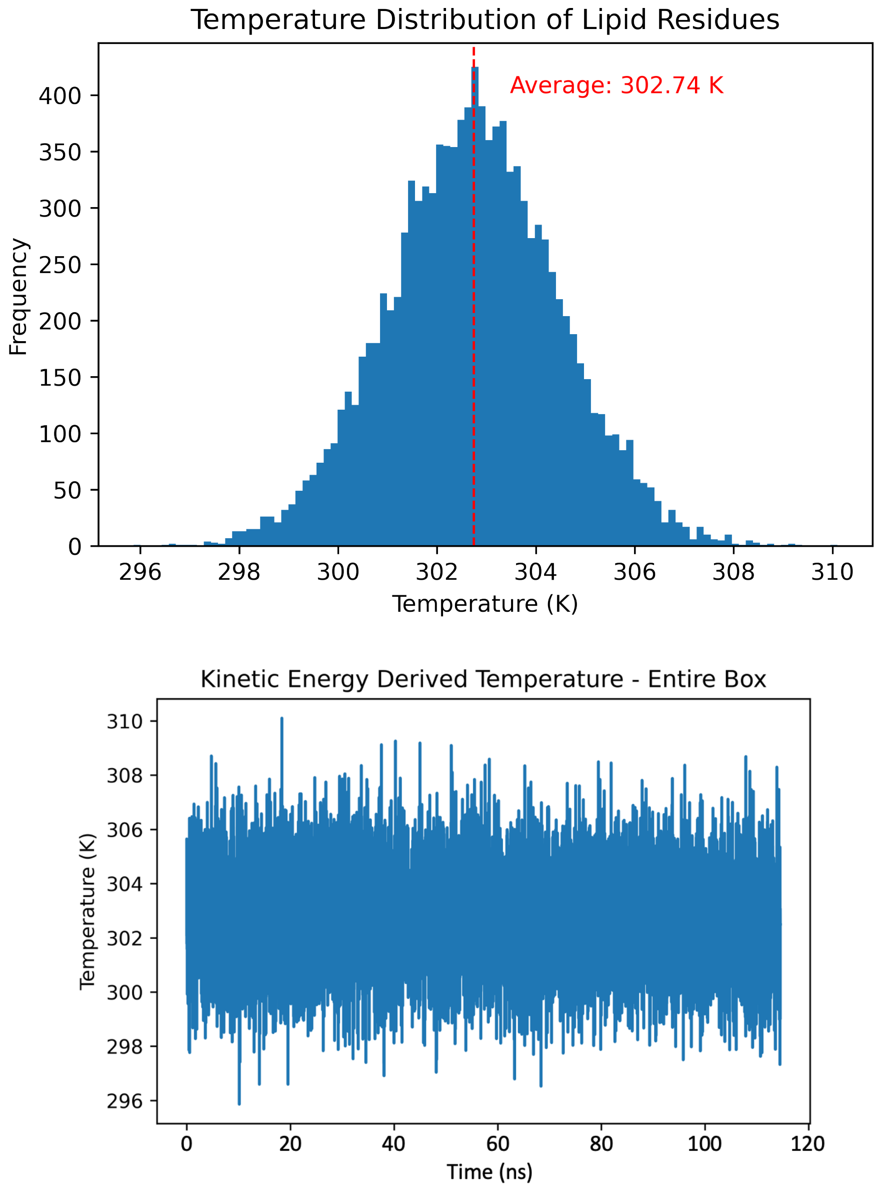


Figure S3: Top: Distribution of kinetic energy-based temperatures, calculated with the equipartition theorem using an MD input temperature of 306 K. Bottom: Time series of the kinetic energy-based temperatures.

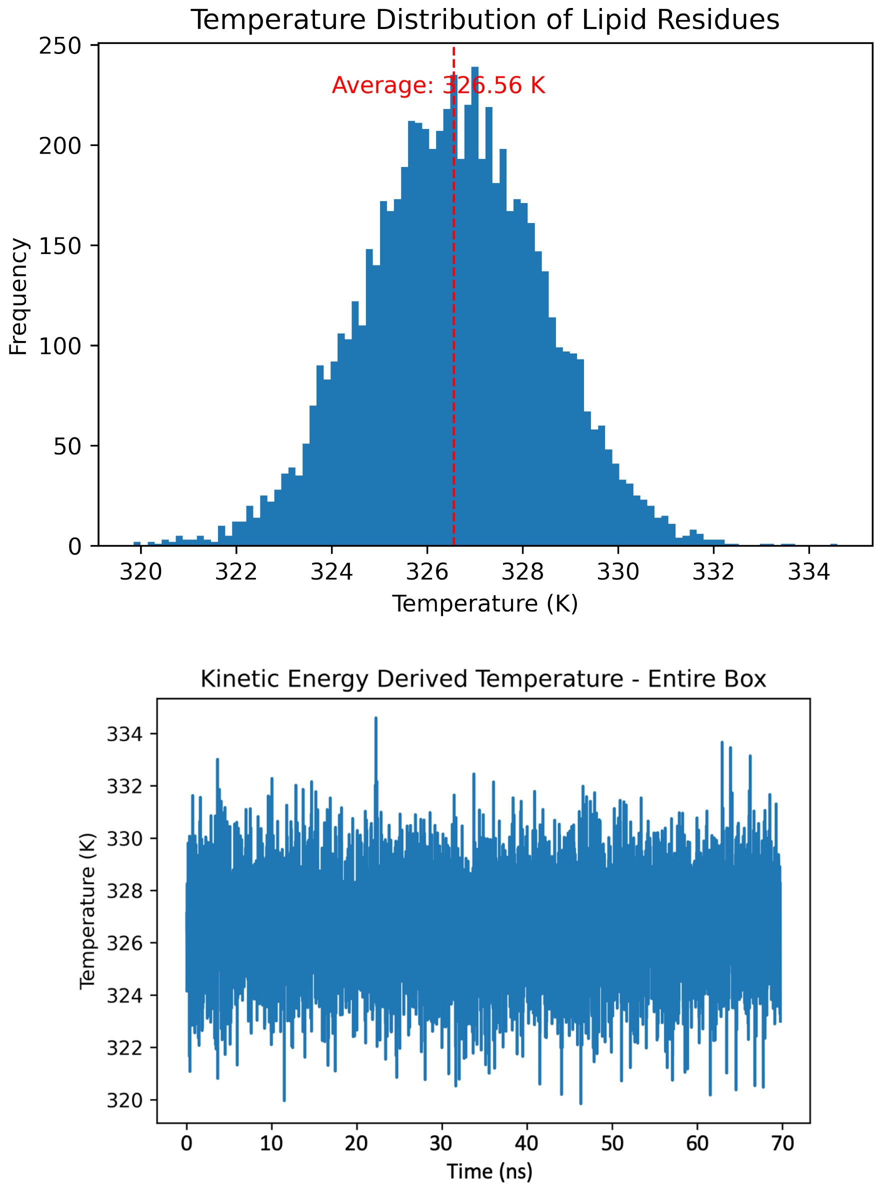


Figure S4: Top: Kinetic energy based calculation of temperature, using equipartition theorem using an MD input temperature of 330 K. Bottom: Time series of kinetic energy based temperature calculation.

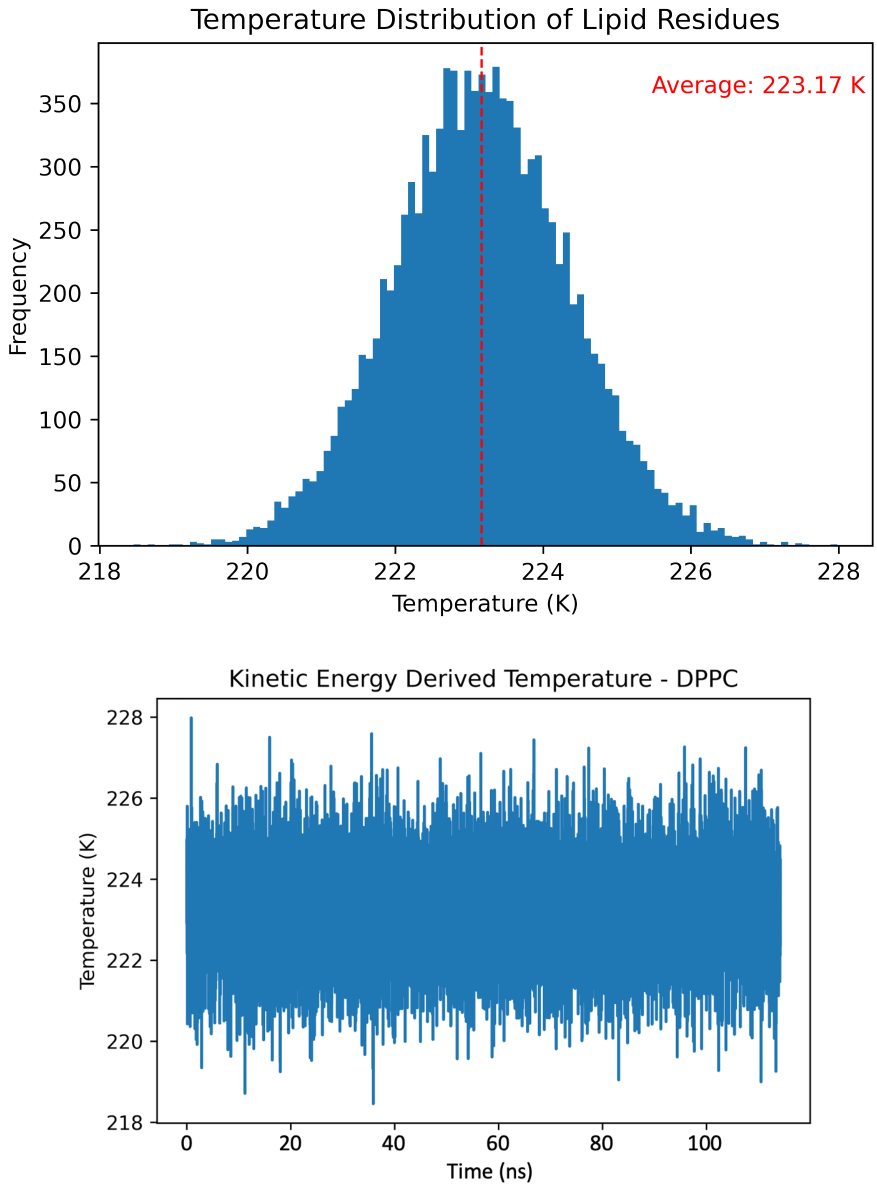


Figure S5: Top: Kinetic energy based calculation of DPPC temperatures, using equipartition theorem using an MD input temperature of 280 K. Bottom: Time series of kinetic energy based temperature calculation.

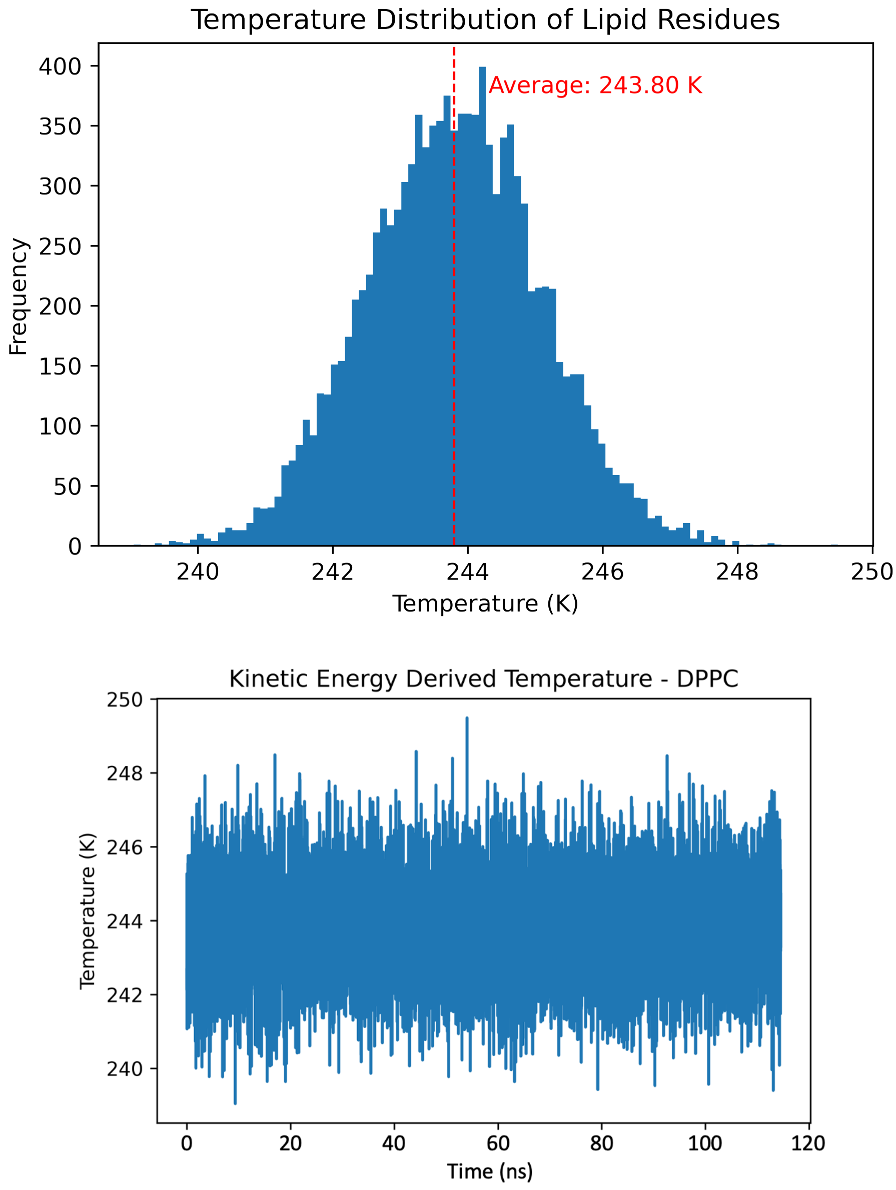


Figure S6: Top: Kinetic energy based calculation of DPPC temperatures, using equipartition theorem using an MD input temperature of 306 K. Bottom: Time series of kinetic energy based temperature calculation.

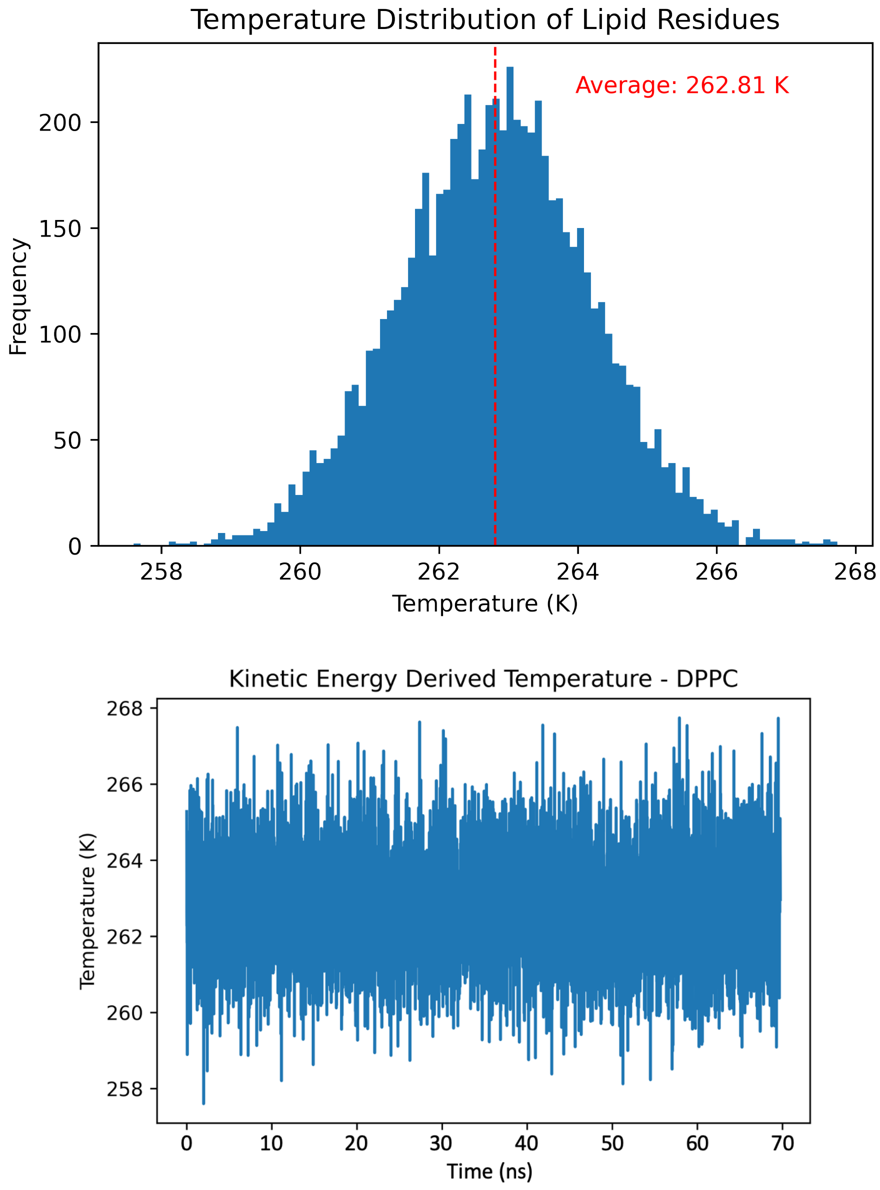


Figure S7: Top: Kinetic energy based calculation of DPPC temperatures, using equipartition theorem using an MD input temperature of 330 K. Bottom: Time series of kinetic energy based temperature calculation.

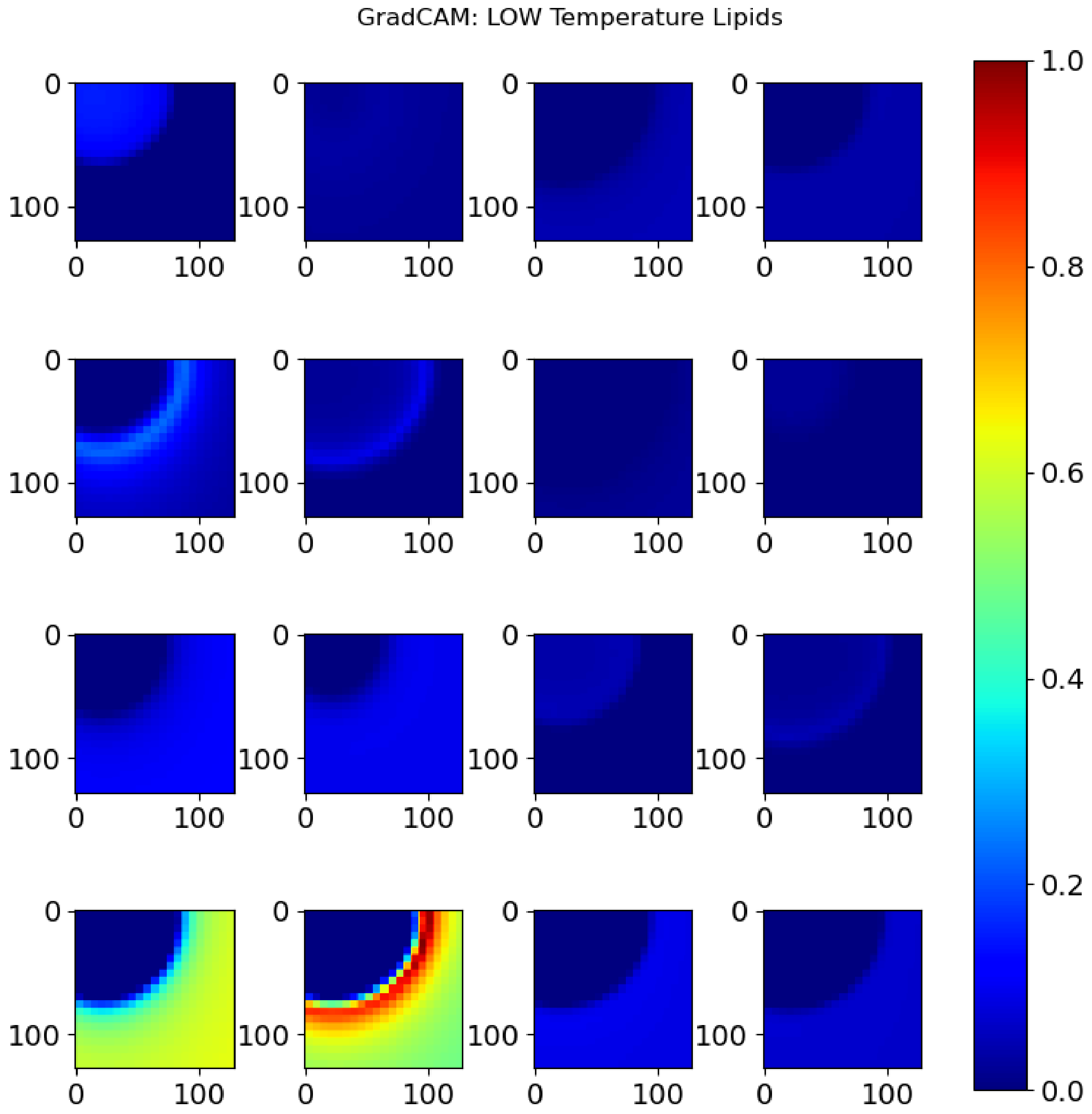


Figure S8: Explainable AI method. The GradCAM method stands for gradient based class activation map which computes hidden layer vector's gradient with regard to predicted class by `MembTDA`. The computed hidden layer vector's gradient is then projected onto a persistence image. For each of the 16 randomly chosen $H_1$ persistence images from low-temperature DPPC lipids (<300 K), we compute corresponding gradient of each lipid's persistence image. The color bar indicates the magnitude of gradient value; however, the exact value does not matter for analysis. For most of the images, it is important that we *find* quarter circle pattern in the top left side of each persistence image. This indicates we have persisting topological features (i.e., existence of hole) captured in ordered/rigid configuration of lipid tails.

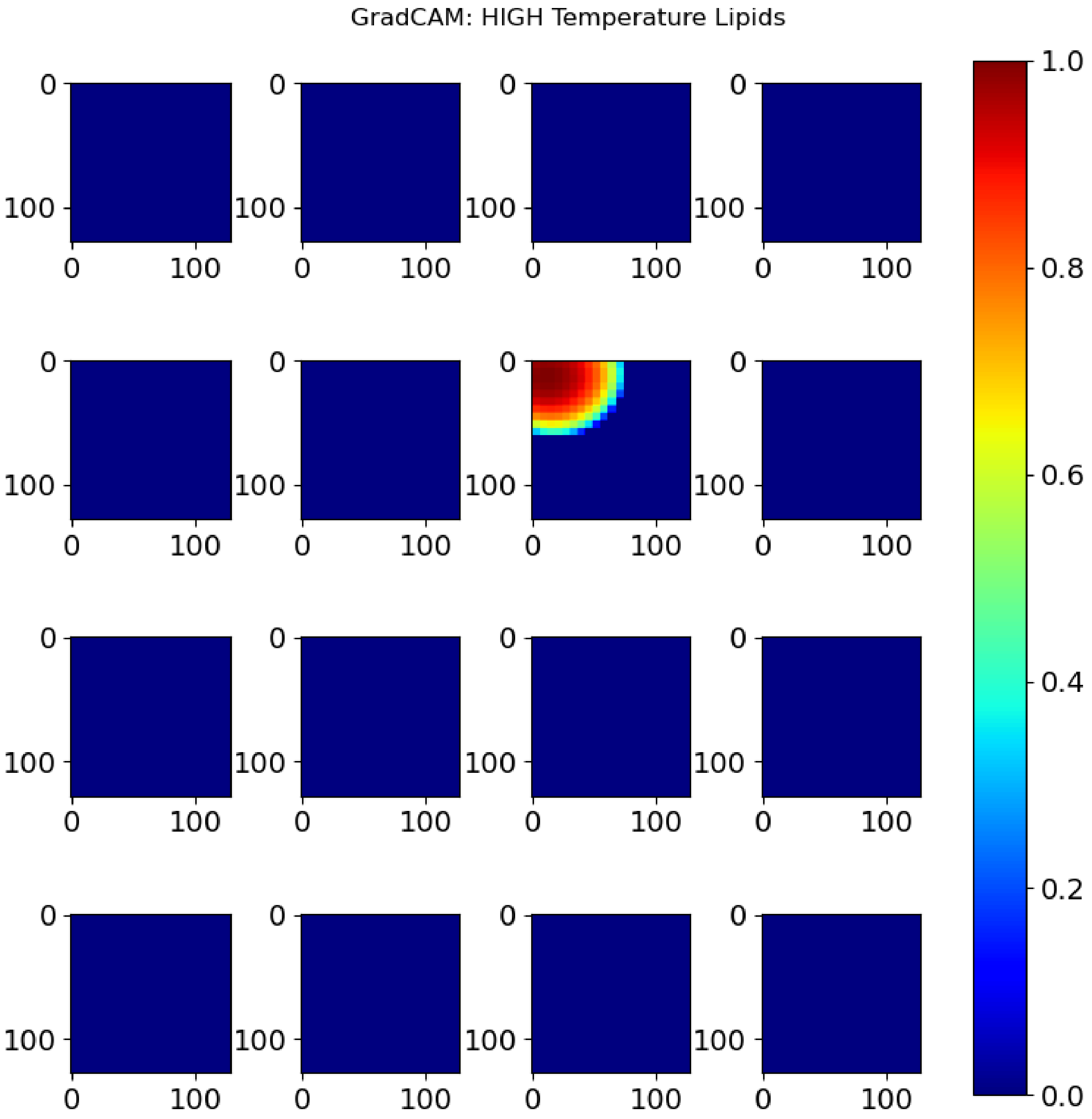


Figure S9: Explainable AI method. The GradCAM method stands for gradient based class activation map which computes hidden layer vector's gradient with regard to predicted class by `MembTDA`. The computed hidden layer vector's gradient is then projected onto a persistence image. For each of the 16 randomly chosen $H_1$ persistence images from high-temperature DPPC lipids (>310 K), we compute corresponding gradient of each lipid's persistence image. The color bar indicates the magnitude of gradient value; however, the exact value does not matter for analysis. For most of the images, it is important that we *do not find* quarter circle pattern in the top left side of each persistence image. This indicates we have lost persisting topological features (i.e., existence of hole) captured in disordered configuration of lipid tails.

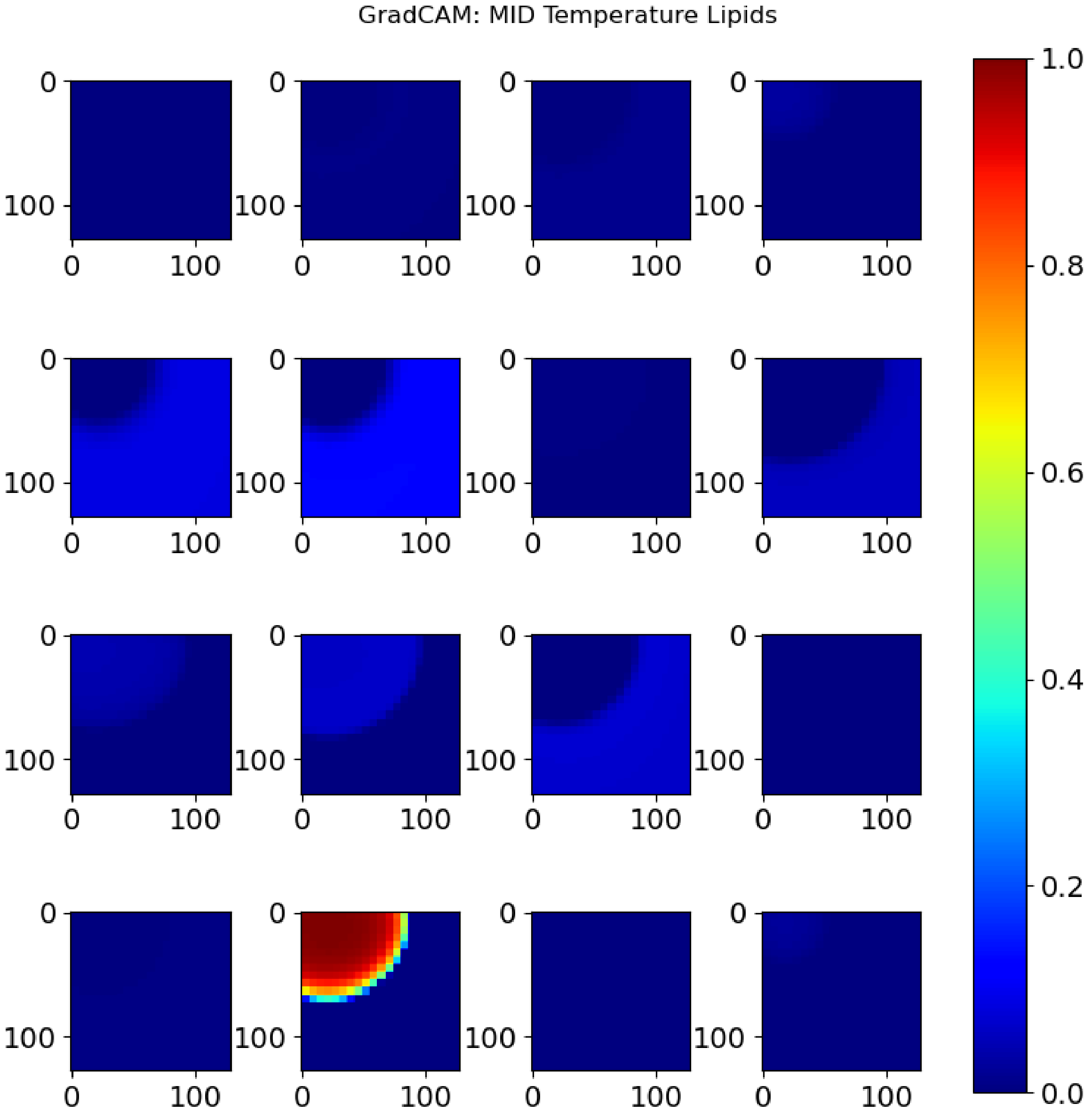


Figure S10: Explainable AI method. The GradCAM method stands for gradient based class activation map which computes hidden layer vector's gradient with regard to predicted class by `MembTDA`. The computed hidden layer vector's gradient is then projected onto a persistence image. For each of the 16 randomly chosen $H_1$ persistence images from melting-temperature DPPC lipids (=306 K), we compute corresponding gradient of each lipid's persistence image. The color bar indicates the magnitude of gradient value; however, the exact value does not matter for analysis. For most of the images, it is important that we find quarter circle pattern in the top left side of each persistence image for *only about half* of the persistence images. This indicates we have persisting topological features (i.e., existence of hole) captured in mixed configuration of lipid tails. That is, at a melting temperature, we expect to find two distinct distributions of both ordered and disordered lipids, as demonstrated here.

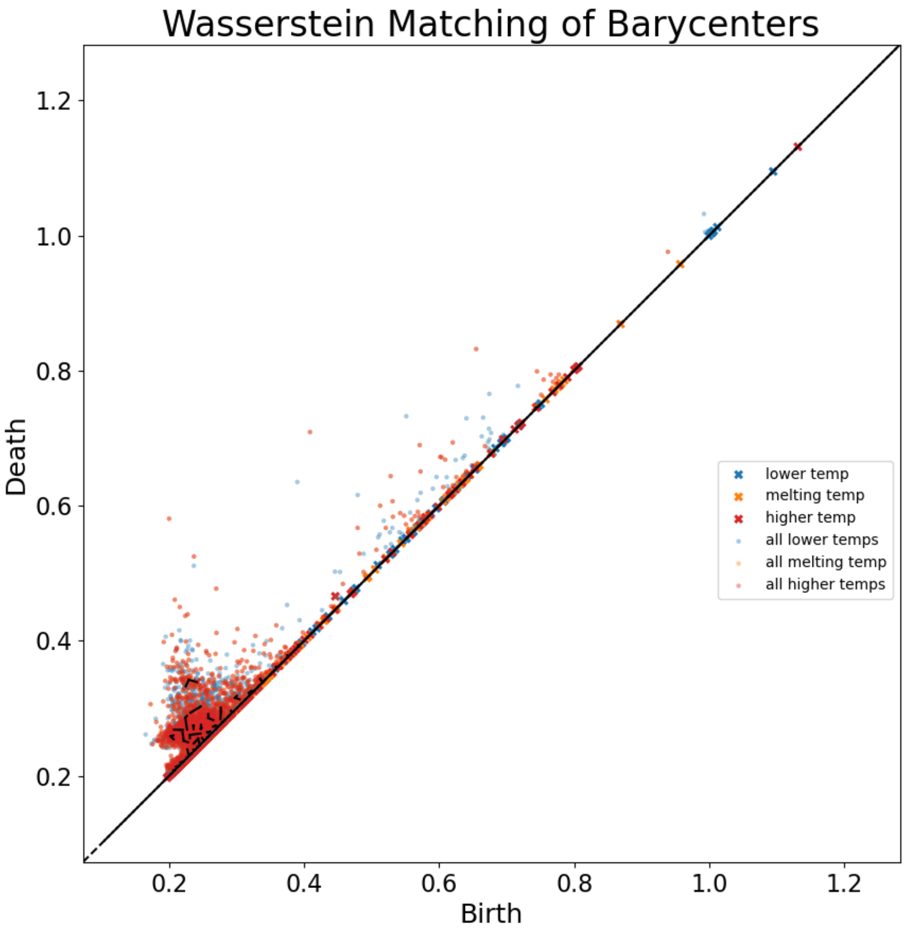


Figure S11: Wassertein matching of all ranges of low-melting-high temperatures. We show the overall $H_1$ Wasserstein matching diagram. We chose 100 random lipids, each from three temperature categories, i.e. low, melting, and high temperatures, resulting in a total of 300 samples, calculated persistence diagrams and overlaid them with colors labeled in the legend (i.e., labeled "all lower temps" with a "**o**" marker; "all melting temps" with a "**o**" marker; "all higher temps" with a "**o**" marker). Then we calculate barycenters of each temperature categories 100 sample persistence diagram birth-death points (i.e., labeled "lower temp" with an "**X**" marker; "melting temp" with an "**X**" marker; "higher temp" with an "**X**" marker). Barycenters are centroids of non-linear systems such as birth-death points of persistence diagrams.[48] In our case, we end up with three barycenter persistent diagrams (i.e., one for each low, melting and high temperature). Additionally, Wasserstein distance is a metric to compare the similarity between two persistence diagrams. The calculation of the Wasserstein distance discerns which points of one diagram are similar to those of another diagram, hence *matching* birth-death points between two barycenter diagrams. These *matching* points are represented as edges connecting two barycenter points between two barycenter diagrams (i.e., an edge ▬ between "**X**" to "**X**", as well as between "**X**" to "**X**"). However, in this diagram, such matching points are not clear because these matching points are concentrated in birth-and-death ranges of 0.2 to 0.4; therefore, we show a close-up view in the following Figure S12.

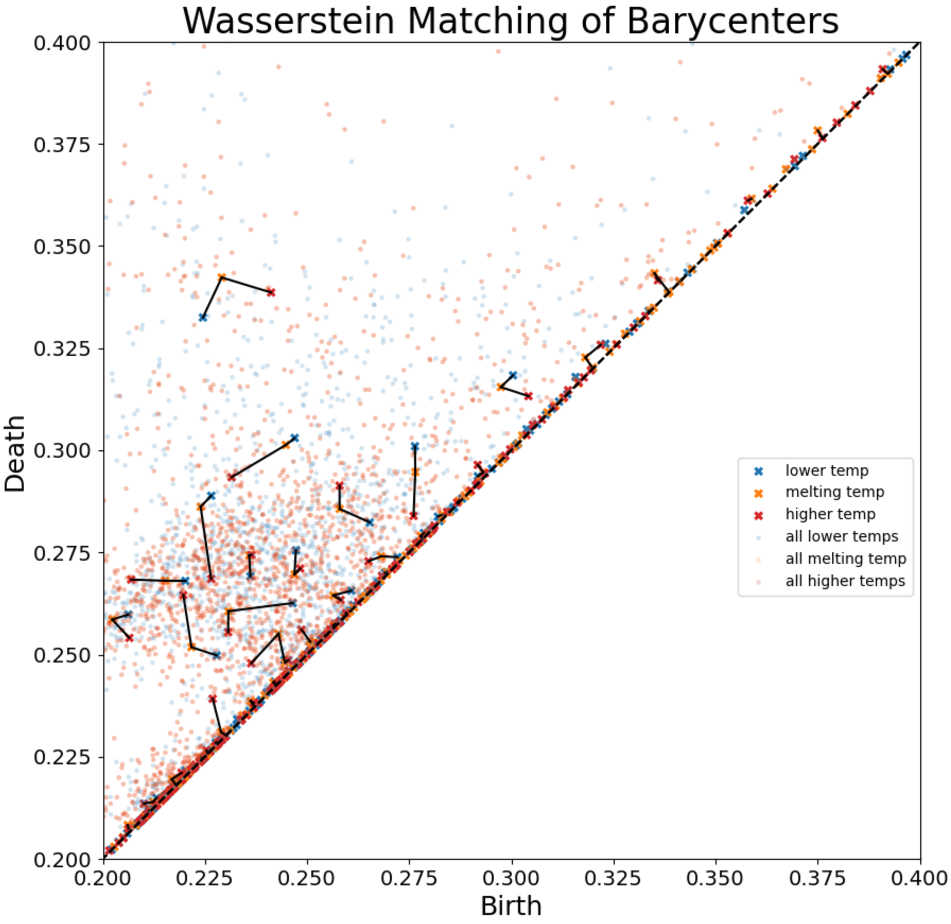


Figure S12: Wassertein matching of specific ranges of low-melting-high temperatures. We show close-up view of the $H_1$ Wasserstein matching diagram in the 0.2-0.4 birth-and-death range for a better visualization. From the Wasserstein diagrams, we can conclude that there are prominent $H_1$ features (holes) appearing and persisting (i.e., death minus birth) about 0.2 to 0.3 $\alpha$ filtration range. In this close-up figure, we can see that the barycenter persistence diagrams from three different temperatures show movement, indicated by edge connections (i.e., an edge ▬ between "**X**" to "**X**", as well as between "**X**" to "**X**"). Although not perfect, we may draw conclusion from the close-up Wasserstein matching diagram that melting temperature barycenter persistence diagram points tend to be situated in-between low and high temperature barycenter points, hence serving as transition points between two lipid phases. This shows that there are distinct persistence data encoded across three lipid temperature categories, indicating temperature dependent encoding of topological features. This is consistent with the idea that `MembTDA` maps out the underlying configurational landscape as seen in Figure 1 where configuration of lipids and temperature are joint probability variables.

#### Deep Learning Model for Processing Persistence Images

In our work, we used `ViT` or related deep learning models such as the well-known convolution-based architecture for prediction of effective temperatures. The `ViT` architecture we used was based on window attention of `Swin Transformer version 2`[25 44] where at different stages, different window sizes are used to aggregate information of the local and neighboring window pixels and patches of a given image. The convolution model we trained with was `ConvNeXt`,[26] which has shown to be comparable with transformer-based models in natural image classification.

We mainly utilized `ViT` architecture since both transformer and convolution models show comparable results within our internal testing. Throughout this study, we refer to `Swin Transformer version 2` as `ViT`. Other architectures experimented with included vanilla `ViT`[24] and `RestV2`,[57] which were not suitable due to CUDA memory errors or high losses.

A vision-based neural network architecture was used for the following reasons: First, it is difficult to directly utilize persistence diagrams or bar codes for training. Second, generating image format input (i.e., persistence images) from raw persistence diagrams or persistence bar codes is a trivial transformation using kernel estimation of each birth-persistence data point. The inherent possibility of variable sized persistence data, mandates feature size agnostic neural network architectures. Persistence diagrams/bar codes from data such as lipid *xyz* coordinates, may produce different sized persistence data for $H_0$ (connected components) and/or $H_1$ features (holes). The size of $H_0/H_1$ features corresponds to how many birth-death pairs are present in the respective homology group. This is why simpler neural network architectures such as multilayer perceptron (`MLP`) or non-neural networks such as decision tree methods cannot be used, as these ML models require fixed feature sizes. Some ways of dealing with variable input sizes include using padding to mask out irrelevant features or flexible adjacency list based graph neural network approach as elaborated in.[22,58] However, the persistence diagram can be quite sensitive to the existence of $H_0$ and $H_1$ features, leading to large noise in the generated birth-death feature. In our internal tests, we confirmed that a graph neural network based approach with `PhysNet`[22,59] as a backbone ML model with persistence diagrams as input could not learn meaningful patterns, yielding high prediction loss.

An image input is easy to generate from persistence diagrams and contains information of the diagram without the need to account for variable feature sizes. This is because persistence images are generated by calculating the statistics, minimum and maximum values, of all the persistence diagram training data. In addition, the persistence images are reshaped into regular grid images and scaled with with respect to the minimum and maximum value statistics. In our work, we scale the persistence image pixel values from $-1$ to 1, with a fixed image size of $128 \times 128$, which is a convention typically used in ML-based image processing. By transforming otherwise less amenable, less regular, and variable-size PH dataset into scaled, regular, and grid-based image input (i.e., persistence images), we can take advantage of powerful and relatively easier to implement computer vision-based neural networks such as `ViT` or `ConvNeXt`, which have shown superior image classification and object detection capabilities.[24,26] In the rest of the article, we refer to our work's neural network as `MembTDA`, unless specified otherwise.

For training, we use homology groups from persitence data of up to only 1 (i.e., $H_1$), since $H_2$ (voids/pockets) or higher groups are expensive to calculate, rarely observed for lipids, and are partially irrelevant since our raw data input consisted of individual lipids, which lack hidden pockets similar to what one might find in proteins. Both $H_0$ and $H_1$ are readily observed in all lipid tails according to our analysis, making finding connected components and holes in a simplicial complex computationally more tractable.

`MembTDA` predicts a temperature class label corresponding to an individual lipid configurational state. This means that given a batch of multiple lipid coordinates, respective persistence diagrams and persistence images are generated; then the generated persistence images are fed to a neural network (i.e., `ViT`-based `MembTDA`), which performs a non-linear transformation to predict corresponding temperature classes in which the lipids were most probably simulated Figure 3. Also, since `MembTDA` predicts a probability for each temperature class, based on which we can also compute an expectation value for the temperature class distribution $\mathbf{E}_{i\in\{280,281,\ldots,330\}} = \Sigma_i[p_\theta(T_i) \times T_i]$, which gives a scalar value that we call an effective temperature, $T_{\mathrm{E}}$. The probability $p_\theta(T_i)$ is the probability of an individual lipid being in a temperature class $T_i$, predicted by `MembTDA`, parameterized by parameters $\theta$. The expected temperature is a weighted average of predicted temperature classes and is what we use to predict a scalar temperature value, $T_{\mathrm{E}}$, for a given lipid configuration.

To train `MembTDA` with persistence images, as described in subsection 2.2, each image is first sliced into multiple patches to allow for batch training. Batch training is a method where a subset of the total dataset (e.g., 128 data points out of 1 million (total) data points from the dataset) is used for optimal training and validation. We iteratively feed batches of data points to an ML algorithm leveraging big data for resource limited hardware. For example, a batch of persistence images comprised of images of $3 \times 128 \times 128$ is transformed into a batch of size $(3 \times 8 \times 8) \times (128/8) \times (128/8)$, which is equivalent to a batch of size $192 \times 16 \times 16$. That is, the original image, which had a $128 \times 128$ shape with 3 channels, is now transformed to an image that has a $16 \times 16$ shape with 192 channels. The reason for this encoding is the $O(N^2)$ spatial complexity of the attention mechanism, where $N$ is the number of image patches (i.e. complexity is reduced from $128^2$ to $16^2$ during this operation). There are further details of what happens after this patching operation such as shifting windows and aggregating information, which are beyond the scope of this work. For details,[25 44] are recommended for reading.

The objective function is the temperature class prediction, $\mathcal{L}_{\mathrm{CE}}(p_\theta(T_i), T_{\mathrm{true}})$, and expected temperature prediction, $\mathcal{L}_{\mathrm{MSE}}(\mathbf{E}_i, T_{\mathrm{true}})$. Where $\mathcal{L}_{\mathrm{CE}}$ is the cross-entropy loss and $\mathcal{L}_{\mathrm{MSE}}$ is the mean squared error (MSE) loss. In practice, we weight the MSE loss term by $\lambda = 0.1$ for stable training to control the magnitude of gradients for each loss: $\mathcal{L}_{\mathrm{total}} = \mathcal{L}_{\mathrm{CE}}(p_\theta(T_i), T_{\mathrm{true}}) + \lambda\mathcal{L}_{\mathrm{MSE}}(\mathbf{E}_i, T_{\mathrm{true}})$. The use of loss functions for both CE and MSE is referred to as bi-objective functions. By training `MembTDA` with bi-objective functions, we can optimize our neural network model for a more robust representation learning of our input. Our overall workflow is described in Figure 3.
